# Predicting CD4 T-cell epitopes based on antigen cleavage, MHCII presentation, and TCR recognition

**DOI:** 10.1101/415661

**Authors:** Dina Schneidman-Duhovny, Natalia Khuri, Guang Qiang Dong, Michael B. Winter, Eric Shifrut, Nir Friedman, Charles S. Craik, Kathleen P. Pratt, Pedro Paz, Fred Aswad, Andrej Sali

## Abstract

Accurate predictions of T-cell epitopes would be useful for designing vaccines, immunotherapies for cancer and autoimmune diseases, and improved protein therapies. The humoral immune response involves uptake of antigens by antigen presenting cells (APCs), APC processing and presentation of peptides on MHC class II (pMHCII), and T-cell receptor (TCR) recognition of pMHCII complexes. Most *in silico* methods predict only peptide-MHCII binding, resulting in significant over-prediction of CD4 T-cell epitopes. We present a method, ITCell, for prediction of T-cell epitopes within an input protein antigen sequence for given MHCII and TCR sequences. The method integrates information about three stages of the immune response pathway: antigen cleavage, MHCII presentation, and TCR recognition. First, antigen cleavage sites are predicted based on the cleavage profiles of cathepsins S, B, and H. Second, for each 12-mer peptide in the antigen sequence we predict whether it will bind to a given MHCII, based on the scores of modeled peptide-MHCII complexes. Third, we predict whether or not any of the top scoring peptide-MHCII complexes can bind to a given TCR, based on the scores of modeled ternary peptide-MHCII-TCR complexes and the distribution of predicted cleavage sites. Our benchmarks consist of epitope predictions generated by this algorithm, checked against 20 peptide-MHCII-TCR crystal structures, as well as epitope predictions for four peptide-MHCII-TCR complexes with known epitopes and TCR sequences but without crystal structures. ITCell successfully identified the correct epitopes as one of the 20 top scoring peptides for 22 of 24 benchmark cases. To validate the method using a clinically relevant application, we utilized five factor VIII-specific TCR sequences from hemophilia A subjects who developed an immune response to factor VIII replacement therapy. The known HLA-DR1-restricted factor VIII epitope was among the six top-scoring factor VIII peptides predicted by ITCall to bind HLA-DR1 and all five TCRs. Our integrative approach is more accurate than current single-stage epitope prediction algorithms applied to the same benchmarks. It is freely available as a web server (http://salilab.org/itcell).

**Author summary:** Knowledge of T-cell epitopes is useful for designing vaccines, improving cancer immunotherapy, studying autoimmune diseases, and engineering protein replacement therapies. Unfortunately, experimental methods for identification of T-cell epitopes are slow, expensive, and not always applicable. Thus, a more accurate computational method for prediction of T-cell epitopes needs to be developed. While the T-cell response to extracellular antigens proceeds through multiple stages, current computational methods rely only on the prediction of peptide binding affinity to an MHCII receptor on antigen presenting cells, resulting in a relatively high number of false-positive predictions of T-cell epitopes within protein antigens. We developed an integrative approach to predict T-cell epitopes that computationally combines information from three stages of the humoral immune response pathway: antigen cleavage, MHCII presentation, and TCR recognition, resulting in an increased accuracy of epitope predictions. This method was applied to predict epitopes within blood coagulation factor VIII (FVIII) that were recognized by TCRs from hemophilia A subjects who developed an anti-FVIII antibody response. The correct epitope was predicted after modeling all possible 12-mer FVIII peptides bound in ternary complexes with the relevant MHCII (HLA-DR1) and each of five experimentally determined FVIII-specific TCR sequences.

## Introduction

Adaptive immunity involves cellular and humoral responses (1). Cellular immunity is primarily mediated by cytotoxic CD8^+^ T cells which recognize peptide antigens presented by Major Histocompatibility Complex class I molecules (MHCI) while humoral immunity requires CD4^+^ T helper cells responding to peptide-MHC class II complexes (pMHCII) to support antibody production by B cells. The humoral immune response pathway proceeds through multiple stages (Figure 1) (2). First, an antigenic protein is endocytosed by antigen presenting cells (APCs). For the MHCII pathway, the protein can then be cleaved in the endosome by acid-dependent proteases into peptides of ∼10-30+ residues. The invariant chain of the MHCII receptor blocks the peptide binding site in the nascent MHCII protein in the endoplasmic reticulum (ER) and facilitates the export of MHCII receptors without peptide ligands from the ER to a vesicle. Next, the vesicle fuses with the endosome that contains the whole antigen or its peptides. The invariant MHCII chain is then cleaved, leaving only a non-covalently bound small fragment (CLIP) that continues to block the MHCII peptide binding site. CLIP is subsequently removed by human leukocyte antigen DM (HLA-DM, with an MHCII-like structure), allowing for subsequent interactions with antigenic peptides in the vesicle. Additional cutting and trimming of peptides can take place after the formation of a stable peptide-MHCII (pMHCII) complex (3). Next, stable pMHCII complexes are presented on the APC surface. If the pMHCII complex is recognized by a T-cell receptor (TCR) on a CD4^+^ T cell, the cell becomes activated, producing helper cytokines that support B-cell activation, differentiation to plasma cells, and finally antibody generation.

**Figure 1:**
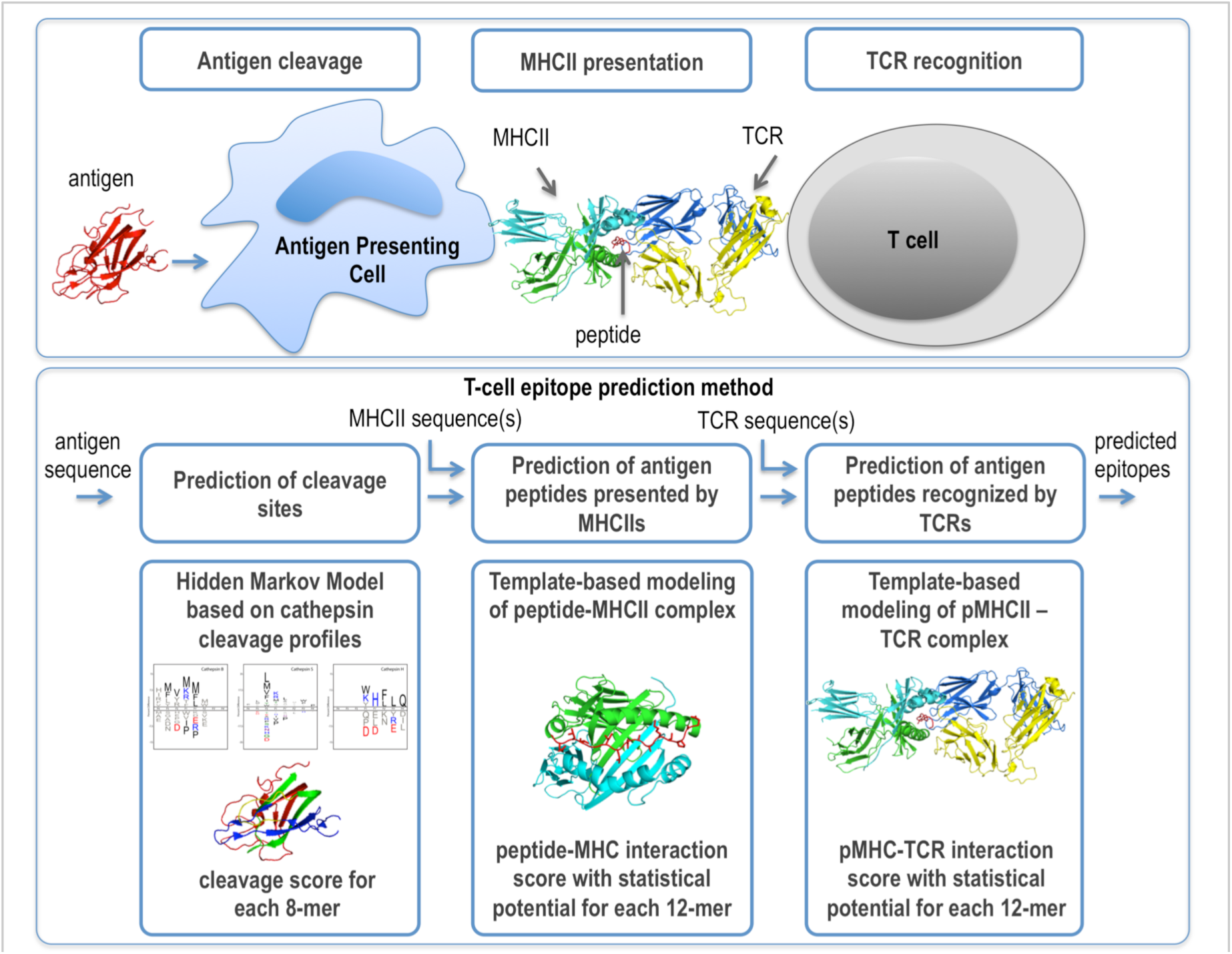
Overview of the 3 steps of antigen processing that are modeled using the current approach.

Here, we focus on predicting T-cell epitopes within an input antigen sequence, restricted to a given human MHCII (Human Leukocyte Antigen, HLA) and to specific TCR sequences. Knowledge of immunodominant epitopes is helpful in designing vaccines (4), improving cancer immunotherapy (5), studies of autoimmune diseases (6, 7), and engineering of protein replacement therapies (8). Unfortunately, most current *in vitro* and *in vivo* experimental methods for predicting immunogenicity in pre-clinical settings are resource intensive and relatively slow (5, 9, 10). Therefore, an accurate, rapid, and inexpensive computational approach for assessment of protein immunogenicity would be valuable.

Current computational methods that are informative about T-cell epitopes predict mainly peptide-MHCII affinity (11). They can be divided into machine learning (12) and *ab initio* approaches (13). Machine learning methods train predictors by relying on a large number of experimentally measured binding affinities for multiple MHCII alleles (14, 15), such as those available in the Immune Epitope Database (IEDB) (16). State-of-the-art predictors have an average area under the receiver-operating characteristic (ROC) curve (AUC) of ∼0.8, as shown by independent validation (14, 15). The drawback of machine learning methods is that sufficient experimental datasets are not always available for an MHCII allele of interest, resulting in lower accuracy (12). In addition, because the length of an MHCII-binding peptide can vary from approximately 10 to 30+ residues, mistakes can be made in registering the 9-mer peptide subsequence in the core of an MHCII binding groove (17, 18). It is also hard to account for “flanking” peptide residues on either side of the core 9-mer, which can modulate binding affinity, and only the NetMHCII method currently takes this into account (14). In contrast, current *ab initio* methods rely on the MHCII structure for predicting the affinity of peptide binding (13). These methods do not require experimentally derived peptide binding datasets and can be applied to any MHCII allele. However, their accuracy (AUC from 0.62 to 0.68) tends to be lower than that of the machine learning methods (13).

While there are methods that consider preprocessing of class I peptides (19, 20), the current computational approaches for class II consider only a single step (*ie*, peptide-MHCII binding) in the multiple steps of the adaptive immune response. However, many peptides that are capable of binding to a given MHCII will not actually be presented on the MHCII of an APC (*eg*, because the peptide is not excised from the antigen during cleavage). Moreover, many peptide-MHCII complexes that are presented will not be recognized by a TCR (*eg*, self-peptides that are constitutively presented on APCs). There are six MHC class II genes in humans (HLA-DPA1, HLA-DPB1, HLA-DQA1, HLA-DQB1, HLA-DRA, and HLA-DRB1) of which all but DRA are polymorphic. While peptide presentation by MHCII is essential for immunogenicity, it is clearly not sufficient, as immunogenicity also requires engagement of antigen-specific TCRs from the diverse T cell repertoire (estimated to contain over 1,000,000 unique sequences (21) with a potential diversity on the order of 10^15^ (22)) generated during T cell development. Therefore, ignoring pMHC-TCR interactions leads to overprediction of potential immunogenicity. Including predicted protease cleavage sites in the antigen sequence and assessing the feasibility of ternary complex formation between pMHCII and the corresponding TCR, in addition to evaluating the compatibility between the peptide and MHCII, could significantly improve the accuracy of T-cell epitope prediction. Here, we propose such a combined approach for predicting epitopes, given the antigen, MHCII, and TCR sequences.

First, we designed an antigen cleavage predictor. The minimal cell-free antigen cleavage system requires cathepsins B, H, and S as well as HLA-DM and MHCII (23). Therefore, we experimentally determined specificity profiles of cathepsins B, H, and S, using a global mass spectrometry-based approach (24). These specificity profiles enabled us to design an accurate antigen cleavage site predictor. Second, we designed a structure-based predictor for peptide-MHCII recognition. Using structural modeling and affinity data for a large number of peptide-MHCII pairs in IEDB, we trained an atomic distance-dependent statistical potential (25) that was optimized for assessing the stability of peptide-MHCII complexes. Finally, we designed a structure-based predictor for pMHCII-TCR recognition, enabled by next generation sequencing platforms that can sequence a large number of short DNA sequences covering the diverse complementarity determining regions (CDRs) of the TCRs (26-28). Our approach is modular, for example cleavage sites prediction to filter the list of peptides is optional. The cleavage sites predictor may be more useful for predicting epitopes in pathogen-derived antigens, which are more sensitive to digestion by cathepsins, compared to auto-antigens (29). Finally, our method can also be used when only partial information, such as TCR β-variable regions only, is available, even though the antigen specificity of the complete TCR depends on both α and β chains.

We begin by presenting the integrative structure-based approach (Figure 1). We benchmark the method using 20 known structures of pMHCII-TCR complexes as well as four pMHCII-TCR complexes with defined HLA and TCR sequences only. Finally, we apply the method to predict a validated HLA-DR1-restricted T-cell epitope in blood coagulation factor VIII (FVIII). FVIII-specific T-cell clones were obtained from two hemophilia A subjects who developed a neutralizing antibody response to therapeutic FVIII infusions. The TCR β-variable regions of these clones were sequenced and the resulting sequences incorporated into the variable regions of the TCR models used in our prediction algorithm.

## Results

### Method

The input is an antigen sequence of interest, an MHCII allele sequence, and a TCR β or α+β variable region sequence. The output is a list of potential epitopes (*ie*, peptide sequences in the antigen that are predicted to form a complex with the given MHCII and TCR) as well as structural models of the pMHCII and pMHCII-TCR complexes. The method proceeds in three steps (Figure 1) (Materials and Methods). First, we predict cleavage sites in the antigen sequence based on cathepsins S, B, and H cleavage profiles. These sites can be used to filter the set of all possible 12-mer peptides for the next step. Second, we build a structural model of the peptide-MHCII complex and assess it with an optimized statistical potential. Third, we assess whether or not any of the top-scoring peptide-MHCII complexes are likely to bind to a given T-cell receptor (TCR), based on the score of the best structural model of the ternary peptide-MHCII-TCR complex and the locations of the predicted cleavage sites.

### Substrate specificity profiles for cathepsins B, H, and S

In this step we predict the cleavage sites in the antigen sequence. In a cell-free (*in vitro*) antigen processing system, cathepsins B, H, and S were found to be sufficient for cleavage of model protein antigens (12, 23). Cathepsin S is the major endoprotease involved in class II antigen processing outside the thymus. Carboxypeptidase (cathepsin B) and aminopeptidase (cathepsin H) activities are important for trimming longer fragments bound to MHCII molecules; they are constitutively expressed in all professional APCs. Therefore, we obtained peptide cleavage datasets (24), constructed specificity matrices for the three cathepsins, and designed a cleavage predictor (Figure 2). Cathepsin S has a preference for hydrophobic residues in the P2 position and for positively charged residues in the P1 position, consistent with the profiling of the P1-P4 positions using a combinatorial library (30). The remaining positions have broad specificity, consistent with the major role of this enzyme in antigen processing. The profiles of cathepsins B and H clearly identify them as a predominant di-carboxypeptidase and a mono-aminopeptidase, respectively. Pathogen-derived epitopes have different processing pathways and are quite sensitive to digestion by cathepsins, compared to auto-antigens that are more resistant to digestion (12). As a result, it is recommended to use the cleavage predictor as an initial step when analyzing pathogen-derived antigens, while it may be prudent to generate predictions for auto-antigens both with and without the cathepsin-cleavage prediction filter.

**Figure 2:**
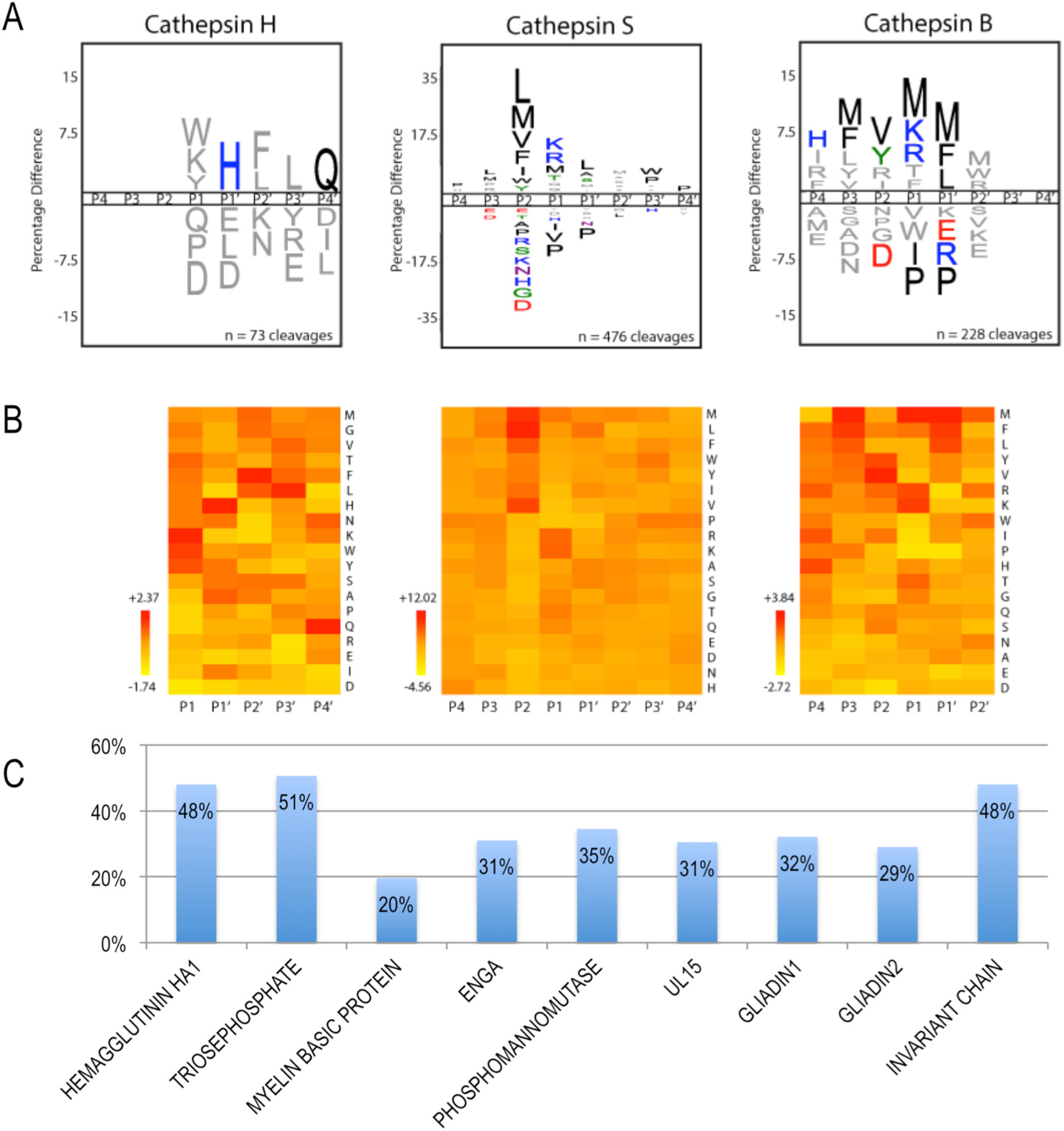
A and B) Cleavage profiles for cathepsins H, S, and B. A) iceLogo representations of substrate specificity. Residues above the line are favored at a given position; residues below the line are disfavored. Statistically significant residues (p ≤ 0.05) are colored according to their physicochemical properties. B) Heat map representations of residue preference clustered and colored by Z-score at each position. Favored residues have Z > 0 and disfavored residues have Z < 0. Cleavage profiles are provided from P1-P4’ for cathepsin H, P4-P4’ for cathepsin S, and P4-P2’ for cathepsin B based on their predominant experimentally-determined cleavage preferences. C) Percentage of 12-mer peptides in the benchmark proteins remaining, following filtering by the cleavage sites predictor.

### Testing on input antigen sequences with known ternary complex structures

To test the accuracy of the method, we used a benchmark dataset of 20 human pMHCII-TCR complexes with crystal structures available in the Protein Data Bank (PDB (31); Table 1). For each complex, we identified and utilized the sequence of the parent protein of the peptide epitope. The benchmark dataset contains antigen sequences from eight proteins (hemagglutinin HA1, triosephosphate isomerase, myelin binding protein (MBP), engA, phosphomannomutase, UL15, and two gliadin proteins). The 20 complexes have some redundancy, since some involved the same MHCII allele, TCR, or peptide epitope sequence. However, the dataset includes 14 non-redundant pMHCII complexes.

**Table 1:**
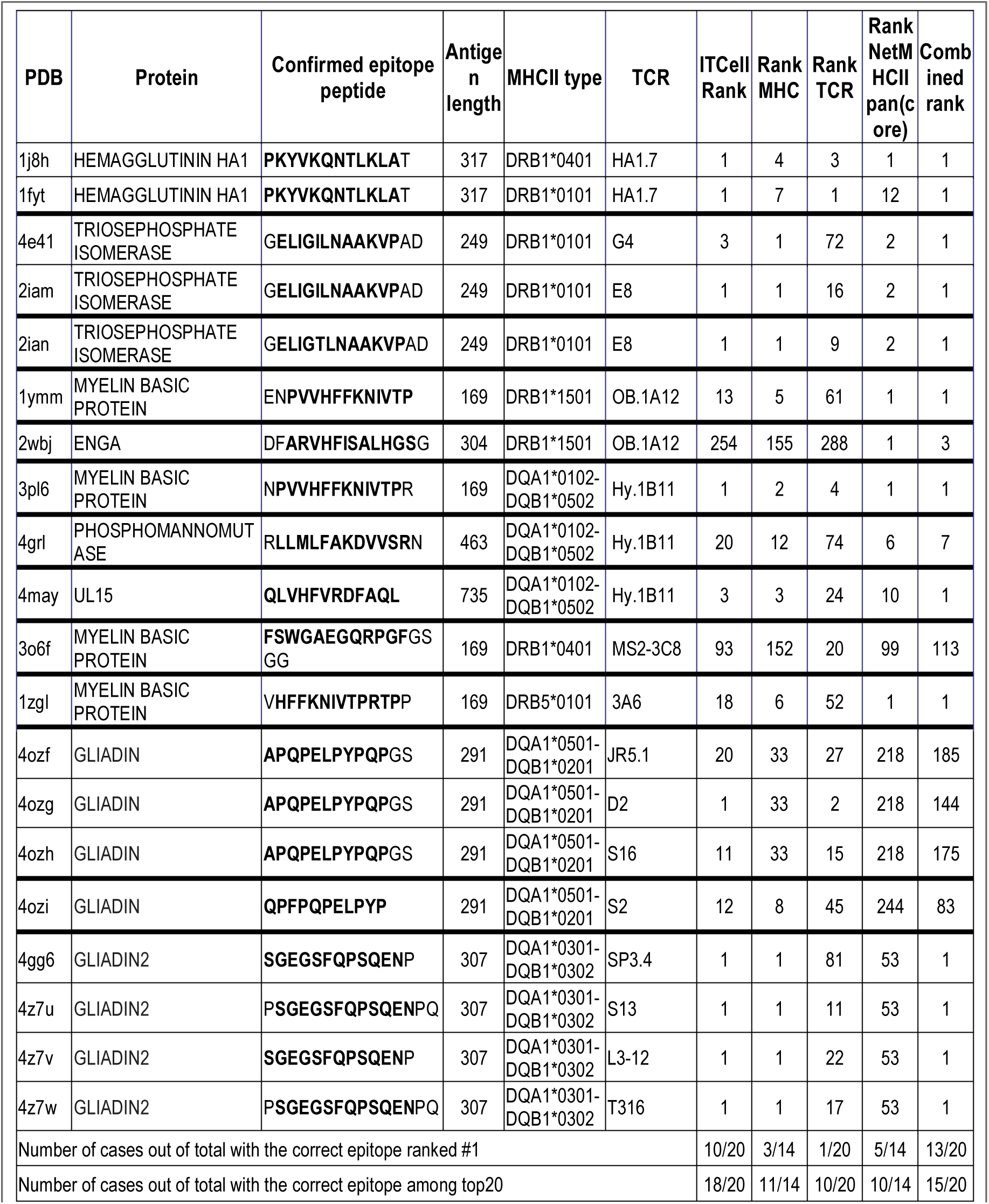
Benchmark of epitope predictions based on crystal structures. The epitope core residues within the peptides are bolded.

For additional benchmarking to test the accuracy of pMHCII complex predictions only, we included an additional 18 pMHCII complexes from the PDB without a known cognate TCR. This combined benchmarking set consists of 32 pMHCII complexes (14 non-redundant pMHCII-TCR complexes + 18 pMHCII complexes).

#### Accuracy of predicting antigen cleavage sites

We applied the cleavage prediction protocol to the sequences of the eight protein antigens in our benchmark set and to the MHCII invariant chain, which is also cleaved by cathepsins in the endosome (Table S2). The method predicted 165 cleavage sites in these nine sequences. In the absence of knowing the true cleavage sites for these nine proteins, we can only check whether or not the predicted cleavage sites lie outside the actual epitopes, as they should. Only 5 of the predicted cleavage sites (three in MBP and two in EngA) lie within the known epitopes. The three MBP epitopes are in fact cleaved in healthy individuals, who thus avoid an immune response, in agreement with the prediction. In contrast, they are not cleaved in patients with multiple sclerosis for whom the epitopes were defined. In these patients, MBP, a short disordered protein, binds to the recycled MHCII molecules without undergoing endosomal cleavage (32). In summary, as expected, the protease cleavage prediction significantly reduced the number of peptide candidates for subsequent testing (by 30% for the benchmark proteins, Figure 2C, Table S2), while eliminating few of the true epitopes.

Additional benchmarking was performed using sequences of 1,000 MHC binding peptides from 280 proteins in the systeMHC dataset (33), which comprise the immunopeptidome of these proteins as determined by mass spectrometry of peptides eluted from antigen-presenting cells. This Hidden Markov Model (HMM)-based cleavage predictor reduced the number of candidate peptide epitopes for MHCII presentation by ∼30%, with only ∼3% of these 1,000 true MHCII binders predicted to be cleaved.

#### Accuracy of predicting pMHCII complexes

Fourteen unique peptide-MHCII pairs from the 20 pMHCII-TCR structures in our benchmark were used for validating prediction of peptide side-chain orientations and peptide registration in the MHCII binding cleft. Input peptide-MHCII templates were taken from the MHCII allele dataset (Table S1) and contained a peptide different from the one that appears in the benchmark pMHCII-TCR complex. Consideration of each peptide was limited to its core 12 amino acid residues, although in principle shorter or longer peptides can be considered by the method. This 12-mer core starts two positions before the traditional 9-mer core peptide register. All possible 12-mer peptides in the antigen sequences were modeled into the complex, followed by an assessment of the resulting 3D model. The rank of the actual epitope peptide was determined and compared to that computed by the *NetMHCIIpan-3.1* predictor (15) (Tables 1 and S3). In three cases, the correct epitope was the top scoring peptide predicted using the SOAP score. In eight additional cases, the correct epitope was among the top 20 scoring peptides. For MBP, the correct peptide-MHCII complex (PDB 3o6f) was ranked in the 152^nd^ position, and thus was not identified correctly. A possible rationalization is provided by the low affinity of this epitope for HLA-DR4 and its “loose” accommodation in the MHCII binding cavity (34). In the remaining two cases, we find the correct peptide was ranked in the top 30 solutions, but with an incorrect register of the peptide core in the MHCII cavity compared to the reference X-ray structure. Similar results were obtained with *NetMHCIIpan-3.0* (Table 1). In summary, ranking of the pMHCII models was insufficient to accurately predict epitopes, although it is clearly an essential step for the overall modeling and scoring including other aspects of the system (below).

We performed further independent benchmarking of peptide-MHCII recognition using PDB structures of pMHCII complexes that do not include the TCR. This enabled us to increase the number of non-redundant test cases from 14 to 32 (Table S3). Here, all possible 12-mer peptides in the antigen sequences were used without filtering of predicted cleaved sequences. On average there were 440 12-mer peptides tested for each known MHCII binding peptide (Table S3). In this benchmarking set the correct pMHCII binding register was the top-ranked prediction for 5 (16%) and 4 (13%) of the 32 benchmark cases evaluated by our statistical potential (SOAP) and by NetMHCIIpan, respectively (Table S3, Figure 3). Because these two scoring approaches are based on different information, we tested whether the combined score could improve prediction accuracy. Indeed, using a combined score (sum of normalized scores), we were able to improve prediction accuracy, resulting in the correct pMHCII register receiving the top score for 9 (28%) of the 32 cases.

**Figure 3:**
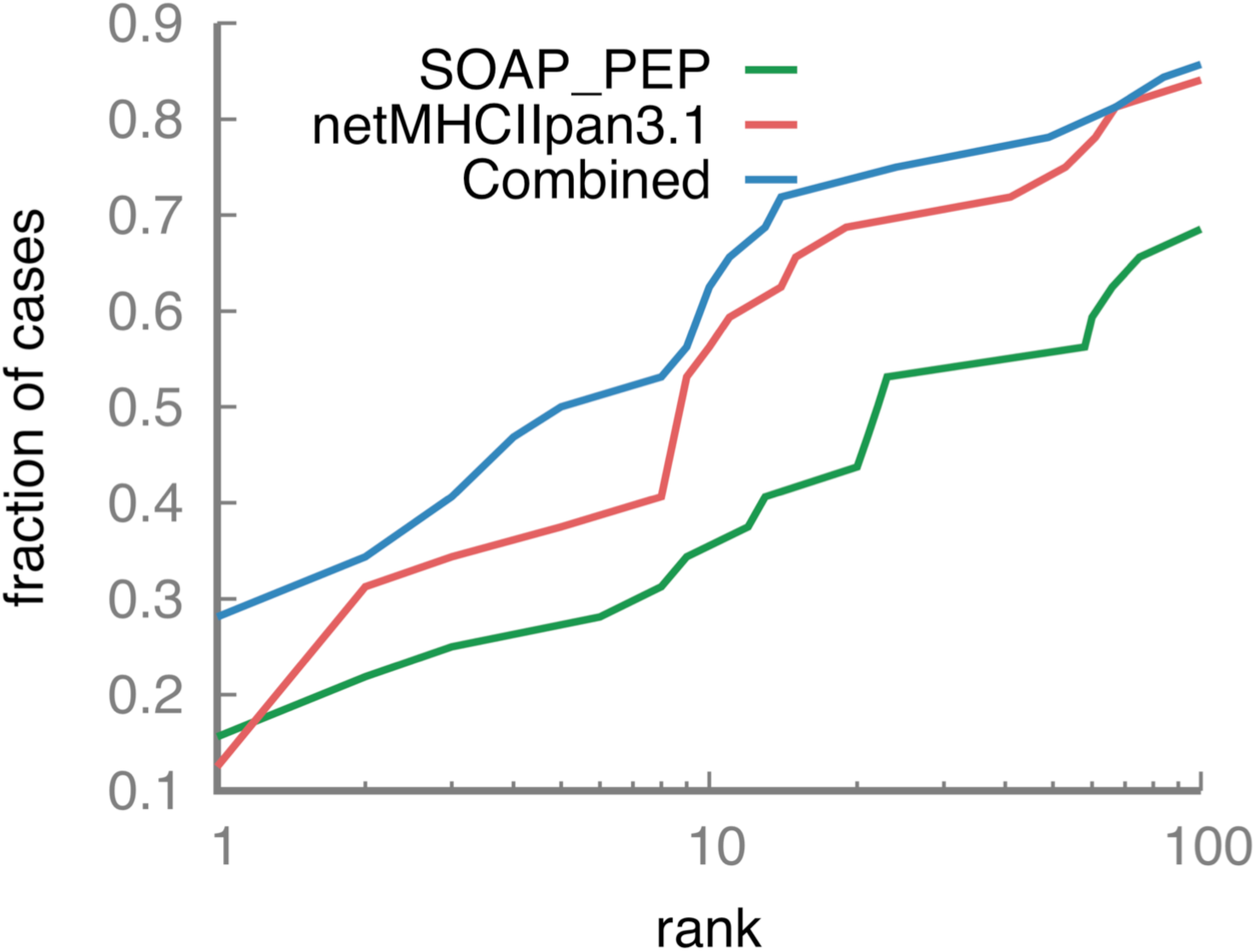
Success rate for SOAP_PEP, NetMHCIIpan3.1 and the two scoring functions combined.

Finally, additional benchmarking to predict peptide-MHCII binding was performed using 29,700 MHCII binding peptides with measured affinities from the IEDB dataset. For each peptide, we identified and utilized the sequence of the parent protein. Our goal was to identify the MHCII binding peptide among all possible 12-mers in the parent protein sequence. Because there are no structures for these pMHCII complexes, their core registers in the MHCII cavities could not be validated. Again, all peptides were used without filtering of predicted cleaved sequences. The correct peptide binding register (overlap of at least 12 peptide residues) was found in the 10 top-scoring peptides for 14% and 22% of the 29,700 benchmark cases using the SOAP statistical potential and NetMHCIIpan, respectively (Figure S1A). The combined score improved the accuracy to 24%. We further checked the performance for peptides with high, intermediate, and low experimentally measured binding affinity. For high affinity peptides, NetMHCIIpan had the best performance: 34% of the cases included the correct MHCII binding peptide among 10 top–scoring peptides (Figure S1B). The combined score improved the accuracy to 36%. For peptides with intermediate affinity, SOAP and NetMHCIIpan had the same accuracy of 15%, while the combined score increased the accuracy to 20% (Figure S1C). For low affinity peptides, the accuracy was 12% and 6% for SOAP and NetMHCIIpan, respectively (Figure S1D). The results show that SOAP, which is a structure-based statistical potential, has the same accuracy over the entire affinity range, while NetMHCIIpan is more accurate for high affinity binders. Moreover, the combined score improves the accuracy for high and medium affinity binders.

#### Accuracy of predicting TCR recognition

We now evaluate whether testing of all possible 12-mer peptides for predicted binding to a known MHCII and a known TCR can identify the correct epitope based on the score of the pMHCII-TCR interface. Similar to the assessment of pMHCII complexes, the correct pMHCII-TCR complex was ranked among the 10 top-scoring models in five (25%) of the 20 benchmark cases (Table 1). For five additional cases, it was among the 20 top-scoring models. Therefore, the scoring of a ternary complex is informative for epitope prediction. Importantly, there was little if any correlation between the accuracy of pMHCII and pMHCII-TCR scoring. As a result, the separate scoring of binary and ternary complexes is complementary and thus increases the accuracy of the integrative approach (below).

#### Accuracy of the entire integrative approach

Our integrative approach ranks potential epitopes based on the pMHCII and pMHCII-TCR scores as benchmarked above (Materials and Methods). In 10 of 20 cases benchmarked against crystal structures of pMHCII-TCR complexes, the correct epitope was ranked as the best by the ITcell scoring function (Table 1). In eight additional cases, the correct peptide was among the 20 top-scoring peptides. The only clearly inaccurate predictions correspond to the MBP (PDB 3o6f) and ENGA epitopes (PDB 2wbj), which have outlying pMHCII scores as discussed above. Combining the ITcell score with the NetMHCIIpan score resulted in 13 out of 20 correct top scoring predictions. However, the combined score for gliadin epitopes was worse, since NetMHCIIpan does not rank these epitopes even among 100 top scoring ones. In summary, the benchmark demonstrates the synergistic contributions of the individual scores, resulting in accurate or nearly accurate prediction of epitopes for all but two cases in the benchmark. Moreover, combining ITcell with NetMHCIIpan further improves the accuracy.

### Testing on sequences without known ternary complex structures

Next, we test the method in a more realistic setting, where the sequences of the antigen, MHCII, and TCR β-chain are known (35), but the corresponding ternary complex crystal structure is not available. In each of these cases, a confirmed MHCII-restricted epitope has been identified by non-crystallographic methods, e.g. MHCII tetramers. Thus, only an assessment of the accuracy of the epitope prediction is performed in this benchmark; an epitope is considered correctly predicted if it overlaps by at least 5 residues with the correct epitope. This second benchmark contains 4 combinations of antigen, MHC, and TCR sequences from mice (Table 2). Because mouse protease cleavage specificities may differ from those of humans, we did not use the APC cleavage predictor in this benchmark. Despite this handicap, the correct epitopes were among the top 11 best scoring peptides in all cases.

**Table 2:**
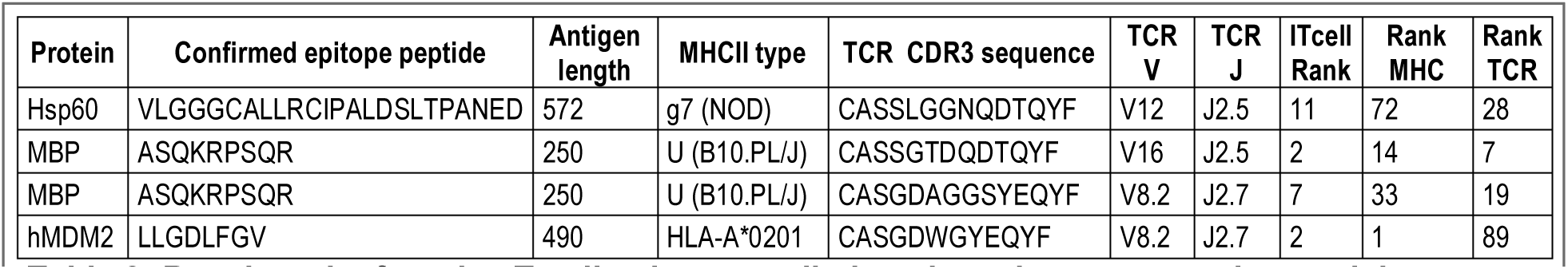
Benchmark of murine T-cell epitope predictions based on comparative models.

### Application to FVIII

To validate the method by applying it to a clinically relevant human immune response, potential *HLA-DRB1*01:01* (abbreviated DR1)-restricted epitopes in blood coagulation FVIII were predicted, using five known FVIII-specific TCRβ sequences as input to the algorithm. TCRβ sequences were determined using cDNA from FVIII-specific T-cell clones that were isolated from two hemophilia A subjects who developed an immune response to FVIII infused to treat their bleeding disorder (36). Epitope mapping using peptide-loaded HLA-DR1 tetramers had identified a 20-mer peptide recognized by these clones as the FVIII peptide spanning residues 2194-2213 (SYFTNMFATWSPSKARLHLQ). The 9-residue HLA-DR1-binding register was subsequently identified experimentally using truncated and sequence-modified peptides (37). The 20-mer peptide and its 10-residue binding register (FTNMFATWSP) were revealed to modelers only after the modeling predictions were shared with experimentalists.

The method was applied in this case without using the cleavage predictor. Based on the benchmark results (Table 1), we expected the actual epitope to be among the 100 top scoring peptides out of 2340 possible overlapping 12-mers. Several 12-mers fully or partially overlapping with the correct 20-mer peptide were among these 100 top scoring peptides. Complexes of the five TCR sequences, the HLA-DR1 receptor, and the core 12-mer (SYFTNMFATWSP) ranked 14, 19, 79, 72, and 52, respectively (Figure 4A-C). This 12-mer was predicted as the most likely epitope for two reasons. First, it was the only consensus 12-mer from the 20-mer peptide since it ranked among the 100 top scoring peptides for all five TCR sequences. Second, it was the only 12-mer fully overlapping with the 20-mer peptide (Figure 4A). Additional 12-mers from the 20-mer peptide were among the 100 best scoring peptides, but they overlapped less (only 5-6 residues) and did not result from considering each of the five TCR sequences. Overall, only six peptides were among the top 100 best scoring for all five TCR sequences; all of them had a Trp residue at position 10 (Figure 4D).

**Figure 4.**
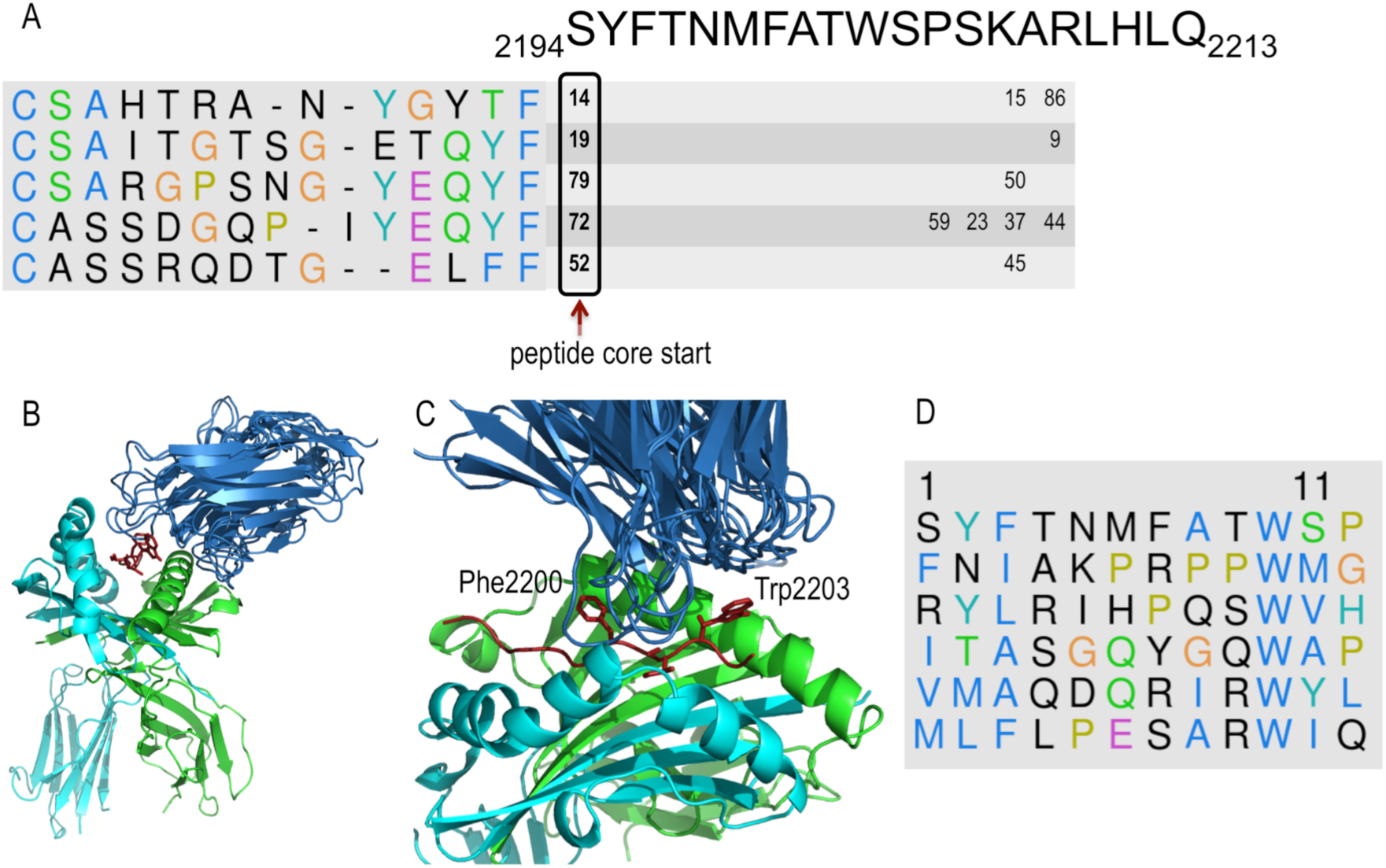
Prediction of FVIII epitopes. A) Consensus based determination of the epitope and its register in the MHCII cavity. CDR3 sequences of the five TCRs and their ranking with respect to different peptide registers in the epitope sequence. Only ranks in the top 100 scores are shown. Peptide FVIII-2194-2205 ranked in the top 100 scoring peptides predicted to bind both HLA-DR1 and each of the 5 indicated TCR-beta variable CDR3 sequences. The predicted consensus core of the epitope for the five TCRs is indicated by an arrow. B) Structural models for the five pMHCII-TCR complexes with the SYFTNMFATWSP 12-mer peptide. The identical MHCII (HLA-DR1) structures within each modeled complex were superimposed, allowing visualization of the different peptide and TCR structural variations. C) The TCR models contact the pMHCII by placing their CDR3 loops in the cavity between FVIII residues Phe2200 and Trp2203. D) The six highest scoring peptides with respect to all five TCRs all have a Trp residue at position 10.

Therefore, using five patient-derived TCR sequences reduced the number of possible 12-residue epitope cores from 2340 to only six FVIII peptides, including the correct epitope core. If we consider consensus peptides only among four of five TCR sequences, we add 12 additional peptides to the prediction list. For comparison, NetMHCIIpan 3.0, which does not rely on TCR sequences, ranks the correct epitope as a weak binder at position 182 only (there is only one NetMHCIIpan prediction per antigen because its predictions do not depend on the TCR sequence).

## Discussion

Epitope prediction is a challenging problem in protein engineering, vaccine design, cancer immunotherapy, and autoimmune disease studies. Even though the immune response pathway involves several steps, most current computational approaches focus on MHCII presentation only. Here, we present an integrative structure-based approach that incorporates additional information by explicitly modeling three steps in the adaptive immune response pathway: antigen cleavage by proteases, MHCII presentation, and TCR recognition. Our approach is *ab initio* as it does not rely on training using known data. The cleavage predictor was built using cleavage sites determined using a random peptide library (24), the peptide-MHC statistical potential was developed using protein-peptide complexes (38), and the pMHCII-TCR statistical potential was developed using a protein-protein docking benchmark (25). We show that the integrative approach is more accurate compared to considering MHCII presentation only (Table 1, Figure 3). For example, the consideration of cleavage prior to MHCII binding can reduce the number of potential peptide candidates by ∼30% (Figure 2C, Table S2).

As for every optimization problem, the accuracy of the method can be limited by the accuracy of sampling and/or scoring. Here, the sampling exhaustively enumerates all possible epitopes in the input sequence(s) and models the structure of the pMHCII-TCR complex. In turn, the scoring function assesses the possible epitopes using the modeled structures. Therefore, the accuracy of the scoring function and the thoroughness of sampling pMHCII-TCR complex structures determine the accuracy of the entire approach.

We addressed the sampling problem by relying on available structures of pMHCII and pMHCII-TCR complexes as structural templates, thereby significantly improving accuracy compared to current *ab initio* prediction of protein-protein complex structures. While the pMHCII complexes do not show significant structural variability (Figure 5A, Cα RMSD <1Å), the pMHCII-TCR complexes show more variability around the canonical orientation (Figure 6A, Cα RMSD < 24Å). To model this variability, we enriched the available structures with additional pMHCII-TCR template structures obtained by protein-protein docking (Figure 5B), resulting in a more uniform distribution of RMSD values (Figure 6C and D, Cα RMSD < 30.0Å, Table S4). To account for TCR loop variability we generated 10 TCR models.

**Figure 5:**
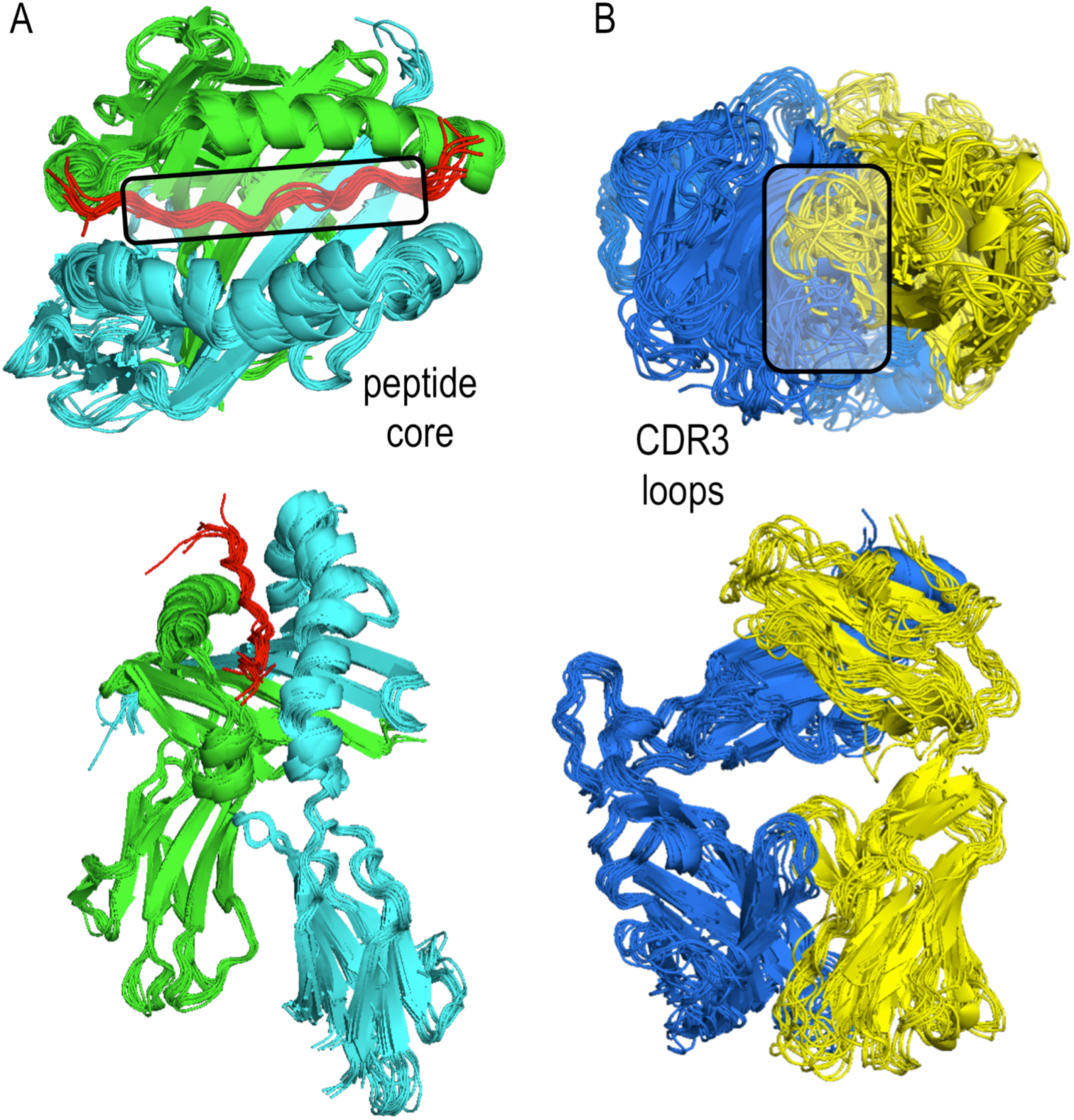
A) Structural alignment of pMHCII complexes from the benchmark set. Alpha and beta chains are green and cyan, respectively. The peptide is shown in red with its core region in the box. B) Structural alignment of TCRs from the benchmark set. TCR alpha and beta chains are shown in yellow and blue, respectively. The variable TCR CDR3 loops are shown in the box. To address TCR loop variability, 10 models are used for each TCR sequence.

**Figure 6:**
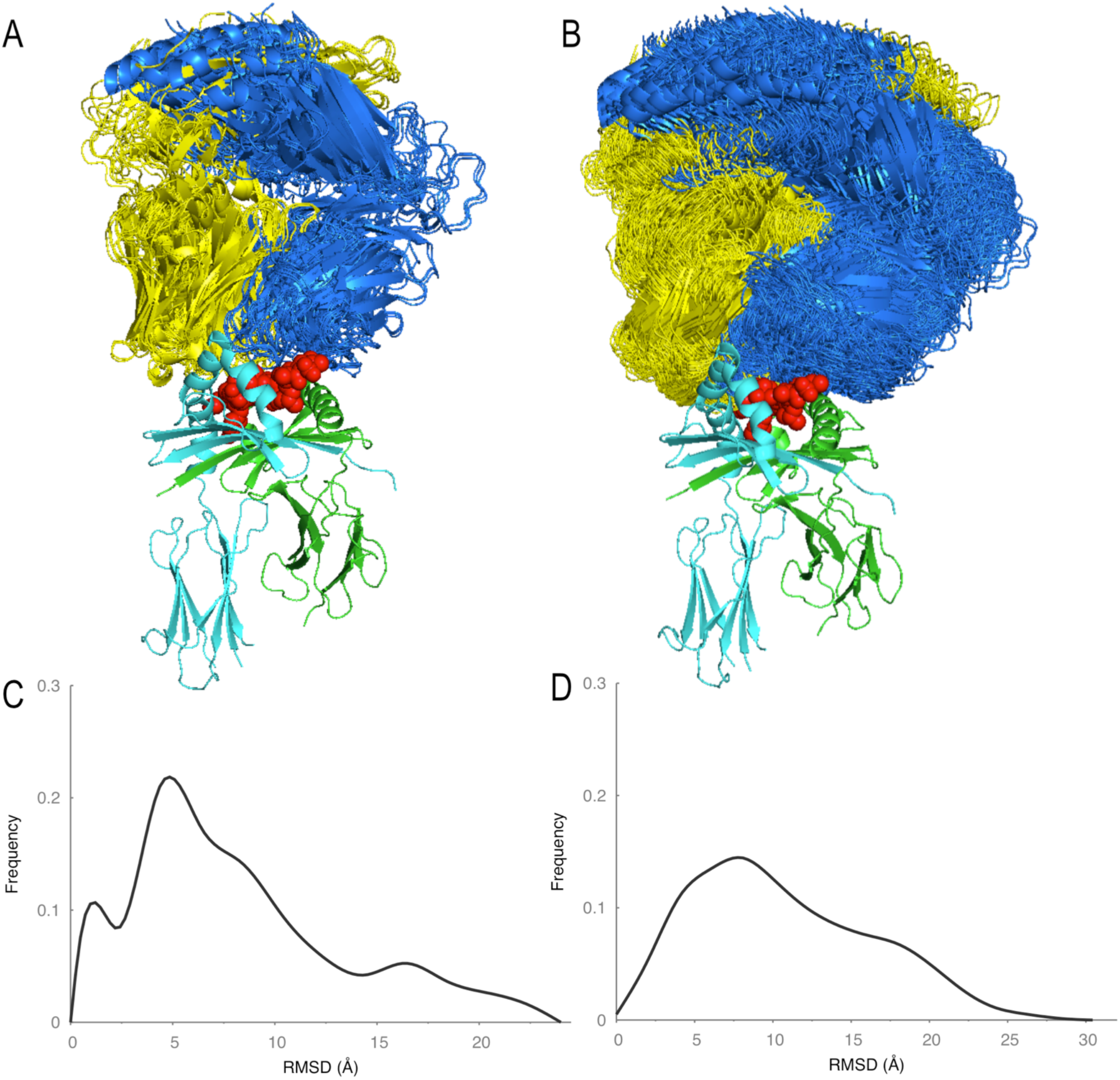
A) Orientations of TCRs with respect to MHCII in the 20 benchmark complexes. B) Orientations of TCRs with respect to MHCII in the 500 template structures generated by docking. C) Distribution of RMSD values between the 20 benchmark pMHCII-TCR structures. D) Distribution of RMSD values between the 500 template structures. Addition of docking generated templates resulted in better coverage of orientation space as indicated by the smoother RMSD distribution on the right.

Our template-based structure modeling approach is not foolproof. For example, recently published structures of an MHCII protein in complex with an insulin peptide and two TCRs revealed a 180° rotation of the TCR compared to the canonical orientation (39). While currently our approach cannot predict this orientation, we could add such novel structures to our template list to include the rotated TCR orientation if it turns out to be common. To maximize scoring accuracy, we used distance and orientation-dependent potentials for peptide-MHC complexes and a distance-dependent potential for pMHC-TCR interfaces. In particular, the *soap_pMHCII* potential was trained for the current problem based on peptides with known binding affinity (Materials and Methods). Statistical potentials have been shown to be useful in a range of protein structure prediction problems (40).

The accuracy of our approach could be further improved in several ways. First, to increase the accuracy of cleavage sites prediction, the current predictor that relies only on experimental cleavage profiles could be upgraded to also consider structural information (41, 42). Moreover, greater coverage of the substrate profiles of the enzymes could be obtained with libraries designed to provide greater representation of amino acid space. Second, pMHCII-TCR recognition accuracy could be improved by modeling the TCR loops in the context of the ternary complex, for example by MODELLER (43). Finally, adding score normalization with respect to peptide length could allow us to predict not only the peptide core, but also the length and conformation of the peptide flanking regions.

Immune responses to protein replacement therapy are a major limitation of protein drugs. Identification and “de-immunization” by removing immunodominant epitopes can prevent immune response and allow wider application of replacement therapy for many conditions, including hemophilia A and B (44), von Willebrand disease (VWD), factor VII deficiency, and Pompe disease (45).

A potential use of our method in the development of biotherapeutics was illustrated by its application to FVIII. Specifically, we predicted a FVIII epitope recognized by five TCRs whose TCR β CDR3 sequences were obtained by multiplex PCR (46) (Adaptive Biotechnologies, Inc., Seattle, WA) of cDNA from FVIII-specific T-cell clones that were isolated from hemophilia A patients who developed an immune response against FVIII replacement therapy (47) (Figure 4). We successfully identified the epitope peptide core among the top 100 choices, among the 2340 possible 12-mer peptides. Consensus scoring for these five TCRs further narrowed the search to six candidate peptides. Moreover, our method has predicted the core of the epitope and its orientation with respect to MHCII and TCR. This strategy could be applied to de-immunizing approaches to identify and eliminate epitopes through strategic amino acid substitutions.

While our method can be applied to any MHC allele without reliance on training datasets like allele-specific machine-learning methods (Introduction), it requires knowledge of the TCR sequences. The gain is the additional information provided by ITCell that associates a TCR with its cognate pMHC. This information could also be used in the context of designing T cells with specific high-affinity TCRs to target known antigens for T cell therapy of cancer or autoimmunity. The method could also aid in identifying the epitopes for lists of TCR sequences that are correlated with a specific condition (such as TCRs of tumor infiltrating T cells, or TCRs found in patients with an autoimmune disease). These TCRs may be identified by high-throughput TCR sequencing (26-28) and determination of the corresponding MHC alleles of the blood donors is straightforward. Moreover, the ITCell method can also be applied to the MHCI pathway (Table 3), combined with existing methods for prediction of proteasomal cleavage and substrate specificity of the transporter associated with antigen processing (TAP). In addition to pMHCI-TCR and pMHCII-TCR, our approach may also be applicable to other immune recognition complexes, such as the pMHCI-KIR3D complexes (48).

## Materials and Methods

The T-cell epitope prediction problem is defined as follows. Given the sequences of an antigen, MHCII allele(s), and antigen-specific TCR(s) that result in the immune response, predict the peptide(s) in the antigen sequence that are both cleaved in the endosome and form a complex with a given MHCII and TCR. The method proceeds in three steps, corresponding to modeling of antigen cleavage, MHCII presentation, and TCR recognition. First, the antigen cleavage sites are identified based on the cleavage profiles of cathepsins S, B, and H. Second, we build a structural model of the peptide-MHCII complex and assess it with a statistical potential. Third, we assess whether or not any of the top-scoring peptide-MHCII complexes can bind to a given T-cell receptor (TCR), based on a score of the best structural model of the ternary peptide-MHCII-TCR complex and the locations of predicted cleavage sites.

### 1. Antigen cleavage

#### Peptide specificity matrices

Peptide specificity datasets for cathepsins S, B, and H were obtained using Multiplex Substrate Profiling by Mass Spectrometry (MSP-MS) (24, 49). We note that in the MSP-MS peptide library, norleucine is used in place of methionine. Recombinant human cathepsin H (catalog #: 7516-CY-010), cathepsin S (catalog #: 1183-CY-010), and cathepsin B (catalog #: 953-CY-010) were purchased from R&D Systems and activated according to the supplier’s instructions. Cathepsin H was activated using bacterial thermolysin (catalog # 3097-ZN-020), which was inhibited with 2 mM phosphoramidon (catalog # EI006) prior to cathepsin H profiling. MSP-MS assays were carried out as described previously (24). Briefly, 0.2 µg/mL cathepsin, a matched no-enzyme control, and a phosphoramidon-inhibited thermolysin control (cathepsin H only) were assayed against a diverse library of 228 tetradecapeptides pooled at 500 nM in D-PBS (pH 6.5) containing 1 mM TCEP. After 15, 60, and 240 min, 30 µL of assay mixture was removed, quenched with 7.5 µL 20% formic acid, and flash-frozen in liquid N_2_. Prior to mass spectrometry acquisition, peptide samples were desalted using Millipore C_18_ ZipTips and rehydrated in 0.2% formic acid. LC-MS/MS data were acquired using a Thermo Scientific LTQ-Orbitrap-XL or LTQ-FT-ICR mass spectrometer that were equipped with a Thermo Scientific EASY-Spray Ion Source, EASY-Spray PepMap C_18_ Column (3 µM, 100 Å), and Waters nanoACQUITY UPLC System. The LC was operated at a 600 nL/min flow rate during sample loading for 14 min and then at a 300 nL/min flow rate for peptide separation over 65 min using a linear gradient from 2% to 50% (vol/vol) acetonitrile in 0.1% formic acid. For MS/MS analysis using the LTQ-Orbitrap-XL, survey scans were recorded over a mass range of 325-1500 *m/z*. Peptide fragmentation was performed using collision-induced dissociation (CID) on the six most intense precursor ions, with a minimum of 1,000 counts, using an isolation width of 2.0 Th, and a minimum normalized collision energy of 25. Instrument parameters for the LTQ-FT-ICR were used as described previously (24). Mass spectrometry peak lists were generated using MSConvert from the ProteoWizard Toolkit (50), and data were searched against the 228-member peptide library using Protein Prospector software (v.5.12.4) (51). All cleavages were allowed in the search by designating no enzyme specificity. The following variable modifications were used: amino acid oxidation (proline, tryptophan, and tyrosine) and N-terminal pyroglutamate conversion from glutamine. Protein Prospector score thresholds were selected with a minimum protein score of 15 and a minimum peptide score of 10. Maximum expectation values of 0.01 and 0.05 were used for protein and peptide matches, respectively. For cathepsins B and S, octapeptides corresponding to P4-P4’ were used as the positive dataset. Octapeptides corresponding to all possible cleavages in the MSP-MS library (n = 2,964) were used as the negative data set. For cathepsin H, selected cleavages were restricted to primary mono-aminopeptidase cleavage products (P1-P4’), due to background thermolysin cleavages, and corresponding pentapeptides were used as the negative dataset (n = 228). iceLogo representations and Z-scores were calculated using iceLogo software (v.1.2) (52). All cleavages identified are available in Supplementary Dataset S2. All raw spectrum (.RAW) files are available at the ProteoSAFe resource (ftp://MSV000079617@massive.ucsd.edu; username: MSV000079617; password: MHCII).

#### Prediction of cleavage sites

The cleavage site predictor was built for each cathepsin individually. First, the MSP-MS results were summarized in a matrix that contained amino acid residue counts from the cleaved peptides for each of the eight positions (P4 to P4’). This matrix was then used to compute a Hidden Markov Model (HMM) for cleavage prediction (53): The residue counts were converted to probabilities, normalized by the natural occurrence of amino acid residues; and the score, *cleavage_score*, is the sum of the log of normalized probabilities. All octapeptides in the antigen sequence are considered for proteolytic cleavage at the P1-P1’ position. For a confident cleavage prediction, we require *cleavage_score* > 3.0, corresponding to at least three times higher likelihood of cleavage than by chance. We first consider cleavage by endopeptidases cathepsins B and S, followed by the aminopeptidase cathepsin H for additional trimming at the N-terminus; we do not use the cathepsin H profile for endoprotease cleavage, because our specificity matrix was constructed using only aminopeptidase cleavage sites.

### 2. MHCII presentation

Given the antigen peptides predicted in the previous step, we build a comparative model of the pMHCII complex for each possible 12-mer peptide. We can rely on comparative modeling due to low *C*_*α*_ RMSD (<1Å) between pMHCII complexes in the PDB (Figure 5A).

#### Human pMHCII template structure dataset

Structures of common human MHCII alleles along with their peptides were extracted from the PDB or modeled using other MHCII structures as templates (Table S1). Because the sequence identity for all MHCII alleles to existing templates in the PDB is high (87-100%), only non-identical side chains were re-packed on a fixed backbone using SCWRL4 (54). Additional alleles can be easily modeled and added to the dataset using the same protocol.

#### pMHCII presentation

Modeling of the pMHCII complex optimizes side chain orientations (54) on a fixed backbone, which is obtained from the template in the human pMHCII template structure dataset. The peptide-MHCII interface in the pMHCII model is scored with an atomic orientation-dependent statistical potential, either *soap_peptide* or *soap_pMHCII,* resulting in the *pMHC_score* for each peptide. These potentials were computed using a previously described procedure (25), as follows: *soap_peptide* was trained using the FlexPepDock dataset of protein-peptide structures and decoys (38); *soap_peptide* was used for modeling pMHCII complexes of known peptide affinity (55); *soap_pMHCII* was trained using the resulting models.

### 3. TCR recognition

### Generation of FVIII-specific T cells and TCRBV sequencing

Two hemophilia A subjects provided written informed consent for use of their samples in accordance with the Principles of Helsinki, under protocols approved by the University of Washington and Seattle Children’s Hospital IRBs. Clones were isolated by single-cell sorting of CD4^+^ T cells stained with phycoerythrin (PE)-labeled HLA-DR1/FVIII2194-2213 MHCII tetramers (Benaroya Research Institute Tetramer Core, Seattle, WA). Clones were expanded and their specificity to HLA-DR1 complexed with FVIII-2194-2213 was confirmed by tetramer staining, proliferation, and cytokine production assays (36). CDNA libraries were constructed and the TCRβ CDR3 regions expressed by the clones were sequenced by Adaptive Biotechnologies (Seattle, WA) as described (47, 56).

#### TCR template structure dataset

Human TCR structures were extracted from the PDB to serve as templates for comparative modeling (Table S5). For each target, 10 models were constructed based on the top 10 templates as ranked by Blast (57), using MODELLER v9.8 protocol for multiple templates (43). To account for the variability of the CDR3 TCR loops (Figure 5B), 10 models were computed. In cases were the TCRα chain sequence was not available, models for TCRβ chain only were constructed and used.

#### pMHCII-TCR template structure dataset

The complex between pMHCII and TCR tends to adopt a canonical orientation (58), but variations are possible. We aligned the 20 complexes in our benchmark dataset based on the MHCII structure and calculated the Cα RMSD between TCRs (Figure 6, Table S4). While most of the structures had RMSD values within 5Å of each other, RMS deviations of up to 24 Å could also be seen (Figure 6C). Therefore, we relied on protein-protein docking algorithm to generate template structures for reliable coverage of possible orientations. We separated the pMHCII and TCR structures from five pMHCII-TCR complexes in the benchmark dataset (PDB codes: 1j8h, 2ian, 2wbj, 4may, and 4gg6) and re-docked them using PatchDock (59, 60). Finally, for each of these 5 complexes we selected the 100 docking models with the lowest RMSD to the crystal structure to serve as templates. The resulting 500 template structures provide good coverage of possible orientations with an average RMSD of 1.75Å (standard deviation 0.7) between the closest template and the crystal structure for the 20 benchmark cases (Table S4).

#### TCR recognition

For each pMHCII model from the previous step, we built a comparative model of the ternary pMHCII-TCR complex, based on each individual ternary complex among the 500 complex templates in the pMHCII-TCR template dataset. The pMHCII and TCR structures were aligned on the template using structural alignment (61). Similarly to the *pMHC_score*, the interface between TCR and pMHCII was then scored with the soap-PP atomic distance-dependent statistical potential derived for protein-protein docking (25), resulting in a *TCR_pMHC_score*. We then selected the best scoring pMHCII-TCR model among the 5,000 computed models (ten TCR models times 500 pMHC-TCR templates).

The final peptide ranking was based on first removing the peptides with a *cleavage_score* >3, followed by ranking the surviving peptides by the sum of *pMHC_score* and *TCR_pMHC_score* (*ie*, the final score). These final scores were converted into normalized Z-scores using the mean and standard deviation of the final scores for all antigen sequence 12-mer peptides.

#### Integration of NetMHCIIpan score

The predicted affinity was converted to a normalized Z-score similarly to *pMHC_score* and *TCR_pMHC_score*. The normalized score was added to other scores.

## Acknowledgments

We thank Dr. Anthony J. O’Donoghue for helpful discussions. Mass spectrometry was performed in collaboration with the UCSF Mass Spectrometry Facility (directed by Alma L. Burlingame). This work was supported by grants from the National Institute of Health (F32CA168150 to M.B.W., R21CA186077 to C.S.C. and A.S., R01 GM083960 to A.S., and P41GM103481 to C.S.C), NHLBI (1RC2 HL101851 and R01 HL130448 to K.P.), Bayer Healthcare Pharmaceuticals (to K.P., D.S. and A.S.).

